# Strong genetic structure and limited connectivity among populations of Clark’s Anemonefish (*Amphiprion clarkii*) in the centre of marine biodiversity

**DOI:** 10.1101/2021.03.17.433695

**Authors:** Hugo Ducret, Janne Timm, Melina Rodríguez Moreno, Filip Huyghe, Marc Kochzius

**Affiliations:** Marine Biology, Ecology and Biodiversity, Vrije Universiteit Brussel (VUB), Pleinlaan 2, 1050 Brussels, Belgium; Molecular Genetics and Biotechnology, Universität Bremen, Bibliothekstraße 1, 28359 Bremen, Germany; Department of Biology, Universidad del Valle, Ciudad Universitaria Meléndez Calle 13 # 100-00, 760000 Cali, Colombia

**Keywords:** Clownfish, Coral Triangle, Indonesia, Phylogeography, Marine conservation

## Abstract

Populations of anemonefish species often show signs of local isolation due limited dispersal potential and oceanographic conditions. Additionally, anthropogenic pressure, such as overharvesting and coral reef exploitation causes reduced population size, eventually leading to local extinction. The understanding of the genetic population structure, as well as the influence of both historical and current connectivity, is required to design effective marine protected area (MPA) networks. In this study, the genetic structure of Clark’s Anemonefish (*Amphiprion clarkii*) populations of the Indo-Malay Archipelago (IMA) is assessed through mitochondrial control region (mtCR) sequences and nuclear microsatellites. Results provided evidence of a significant genetic structure (mtCR: Φ_st_ = 0.42039, Φ_ct_ = 0.63852; microsatellites: F_st_ = 0.01449, F_ct_ = 0.05199). Genetic breaks were identified among Western (Padang Karimunjawa), Central (Sulawesi, Borneo, Bali, Komodo, Timor), and Eastern (Biak) IMA populations, which matches with patterns obtained for congeneric and other coral reef taxa. Due to the restricted connectivity among these three regions, it is suggested to consider them as separate management areas in the design of MPA networks.

## INTRODUCTION

The anemonefish *Amphiprion clarkii* is a small reef fish species of the family Pomacentridae, commonly encountered in tropical shallow waters of the Indo-Pacific. Though juvenile individuals settle in coastal waters onto host sea anemones, larvae are planktonic and their movement can be strongly influenced by sea surface currents (SSC), allowing them to disperse to other areas (Simpson et al. 2014). The pelagic larval duration (PLD) is rather short for *A. clarkii* (16-24 d, Ye et al. 2011), suggesting restricted dispersal potential. This notion is supported by several studies reporting very high levels of self-recruitment in anemonefish (*Amphiprion chrysopterus* 27%, Beldade et al. 2012; *A. perideraion* 46.9%, *A. ocellaris* 48%, Madduppa et al. 2014) that varies among species, location and season (Timm et al. 2012). These ecological features are likely to lead to locally isolated populations exchanging very few migrants, which are prone to suffer from reduced population size. This already happened as high fishing pressure is put upon these species (Madduppa et al. 2014; 2018). Anemonefishes, such as Clark’s anemonefish, are indeed popular in the aquarium trade, especially since the release of the animation movies “Finding Nemo” and “Finding Dory”. If too many individuals are harvested at a certain location, and no migration from other populations occurs, the population eventually disappears. These threats increase the need of understanding the patterns shaping the dynamic of these species and designing proper Marine Protected Area (MPA) networks.

The recent development of molecular ecology tools allows to understand the patterns shaping the genetic diversity of several *Amphiprion* species, particularly within the Indo-Malay Archipelago (IMA). Indeed, population genetic studies have already identified several genetic breaks shared by two anemonefish species within the IMA (*A. ocellaris:* Timm and Kochzius 2008; Timm et. al, 2012; *A. perideraion*: Dohna et. al, 2015), but also other coral reef taxa (giant clams: Kochzius and Nuryanto 2008; Nuryanto and Kochzius 2009; Hui et al. 2016, 2017; corals: Knittweiss et al. 2009; van der Ven et al. 2021; sea star: Kochzius et al. 2009). These barriers distinguish several clusters of populations from each other, based on their genetic structure. Most of the studies revealed the existence of two genetic breaks across the IMA, separating three groups of populations from each other. The first one comprises populations from the Western IMA (Eastern Indian Ocean, Java Sea), while the second one includes populations from the Central IMA (Sulawesi Sea, Makassar Strait, Flores Sea, Banda Sea, Maluku Sea and Sawu Sea) and the third one concerns populations from the Eastern IMA (Seram Sea and Western Pacific). However, the genetic population structure of *A. clarkii* remains unknown in this region. No study yet established if genetic breaks are occurring for this species across the IMA. The ecological properties of each species often lead to relatively different differentiation patterns (Timm and Kochzius, 2008, Dawson, 2014, Dawson et al. 2014, Dohna et al. 2015). More information about the genetic structure of *A. clarkia* is needed in order to apply effective conservation plans adapted to this species.

The aim of this study was to assess the genetic structure of *A. clarkii* across the IMA using genetic markers. As ecological differences between sister species might lead to distinct differentiation patterns, a different pattern of genetic structure was expected for Clark’s anemonefish compared to its congeners as a result of its longer PLD (Ye et al. 2011; Dawson, 2014). However, as for its congeners, it was assumed to find highly structured populations and a high genetic diversity in accordance with the life cycle of this demersal species with substrate-attached eggs and limited pelagic larval dispersal. The conservation of endangered species with such life histories requires in-depth studies providing explicit information regarding genetic structure. Therefore, this study will aim at examining the genetic differentiation occurring between populations of Clark’s anemonefish and identify potential genetic breaks among them. This will reveal the location of diversity hotspots for this species and provide the required information for its conservation and the design of management plans.

## MATERIALS AND METHODS

### DNA extraction and amplification

209 fin clip samples of Clark’s anemonefish were collected at 16 stations across the IMA using SCUBA diving, and stored in 96 % ethanol at 4 °C. Genomic DNA was extracted with a commercial kit (peqGOLD Tissue DNA Mini Kit, Peqlab, Erlangen) following the manufacturer’s instructions. A 420-bp fragment of the D-loop segment of the mitochondrial control region (mtCR) was amplified with the primers CR-A (TTC CAC CTC TAA CTC CCA AAG CTA G) and CR-E (CCT GAA GTA GGA ACC AGA TG) (Lee et al. 1995) for all the individuals in the dataset, following a standard protocol detailed in Timm and Kochzius (2008): The 25-μL reaction volume of the PCRs contained 2.5 μL 10× PCR buffer, 0.075 μmol Mg^2+^, 0.25 μmol dNTP mix, 10 pmol of each primer, 0.5 U Taq Polymerase and 10–30 ng genomic DNA of each sample. The temperature profile of the PCR was 95 °C for 2 min, followed by 35 cycles of 95 °C for 30 s, 50 °C for 30 s and 72 °C for 60 s, and the terminal elongation at 72 °C took 2 min. Finally, PCR products were purified with the QIA quick PCR purification kit (Qiagen), and both strands were sequenced on an ABI Automated Sequencers (Applied Biosystems) using the PCR primers.

Eight polymorphic microsatellite loci were amplified for all the 209 individuals at each location. Primers were either FAM- or HEX-labelled and the PCR protocol used by Timm et al. (2012) was also used in the current study (Table 1): the reaction volume of 10 μl contained 1 μl 10x PCR buffer, 0.0125 μmol Mg^2+^, 0.002 μmol dNTP mix, 5 pmol of each primer, 0.1 U Taq polymerase and 10–20 ng genomic DNA of each sample for each locus. The thermal profile was as following: 94°C for 2 min, followed by 35 cycles of 94°C for 45 s, the primer specific annealing temperature (Table 1) for 45 s, and 72°C for 60 s, with a 2-min long terminal elongation at 72°C. Fragment lengths were subsequently measured with an ABI Automated Sequencers (Applied Biosystems) and analysed with the software Genemapper (Applied Biosystems).

**Table 1:**
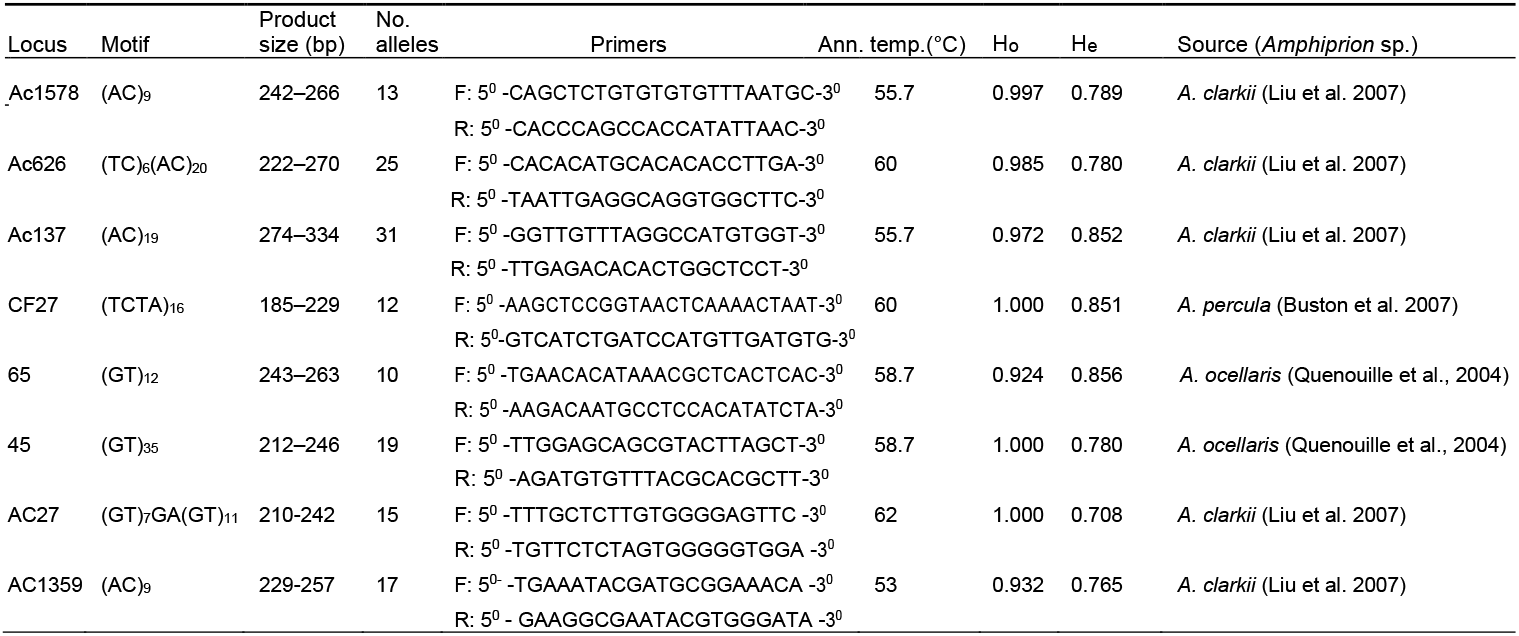
Polymorphic microsatellites loci used for the analysis of the clownfish *Amphiprion clarkii*. Observed (Ho) and expected (He) heterozygosities are indicated for each locus. Ann. Temp: annealing temperature for PCR.

### Control region

Neutrality of the mtCR marker was evaluated to ensure suitability for population genetic analyses. Tajima’s D (Tajima, 1989) and Fu’s F_s_ (Fu, 1997) were used to detect recent population expansion or bottlenecks.

Unless otherwise stated, all following analyses were carried out with Arlequin (ver. 3.5). For mtCR, nucleotide and haplotype diversities were assessed for each sample site (Nei 1987). The overall genetic population structure (Φ_st_), as well as pairwise population differentiation (pairwise Φ_st_), was established through an Analysis of Molecular Variance (AMOVA). The matching P-values obtained with computations were adjusted regarding the false discovery rate (FDR) following Benjamini and Hochberg’s procedure (1995) (multtest, R package 2.9.0). Groups for hierarchical AMOVA testing were chosen to depict regional clustering based on population differentiation, which significance was obtained according to a p-value. Different scenarios were tested according to pairwise Φ_st_-values obtained with the AMOVA.

A haplotype network was constructed from 209 mtCR DNA sequences with the software TCS (v.1.2.1; Clement *et al*. 2000). All haplotypes were included in the haplotype network, from which haplogroups were identified. They were defined as containing less mutational steps within than among them, and single outlier haplotypes were not defined as haplogroups as their position in the haplotype network is questionable (Dohna et al. 2015). The graphical output shown in Fig. 1a was drawn by hand on the basis of what was obtained with TCS. Pie charts show the relative frequency of each haplogroup in each population in Fig. 1b.

**Fig. 1:**
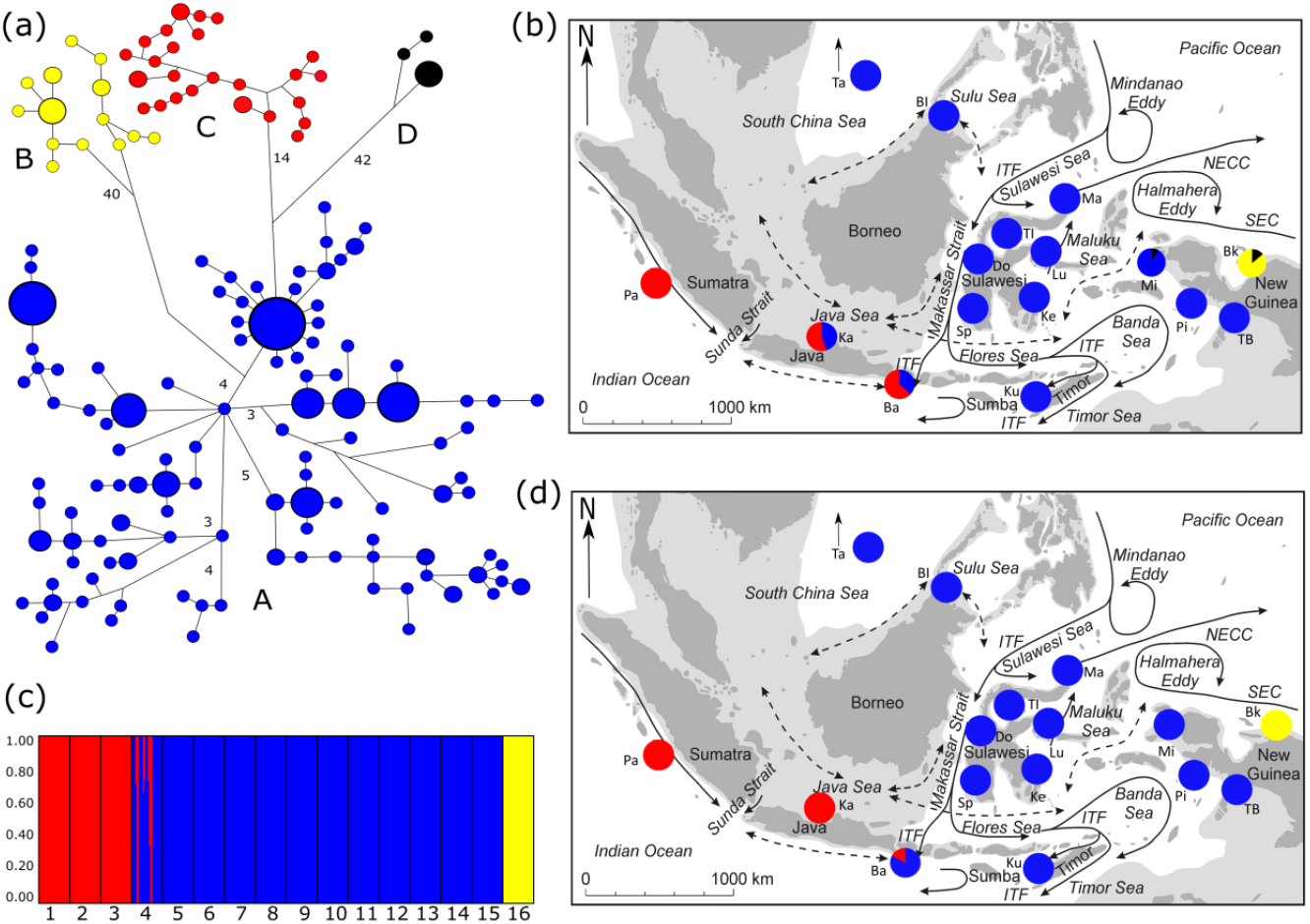
(a) Hayplotype network of *Amphiprion clarkii* based on 209 mtCR DNA sequences. The size of the coloured circles is relative to the number of individuals represented by that haplotype. The smallest coloured circles represent one individual, while the biggest represent eight individuals. Connecting lines without numbers mean one mutational step between haplotypes, and additional mutations are indicated by numbers. (b) Map showing the distribution of the haplogroups identified throughout the populations of the Indo-Malay Archipelago (IMA). (c) Bar plot calculated with the software STRUCTURE for all of the 16 populations (1 = Padang, 2 = Karimunjawa, 3 = Bali, 4 = Spermonde, 5 = Donggala, 6 = Banggi Islands, 7 = Taiwan, 8 = Togian Islands, 9 = Manado, 10 = Luwuk, 11 = Kupang, 12 = Kendari, 13 = Misol, 14 = Pisang, 15 = Triton Bay, 16 = Biak), with K = 3. (d) Map showing the distribution of the genotype clusters identified with STRUCTURE for the populations of the IMA. For both maps, arrows indicate major surface currents, dashed arrows depict seasonally reversing currents, dark grey areas are present-day land formations, and light grey shading indicates marine habitat exposed during the Pleistocene glacial maxima, which led to a 120 m drop in sea level (Voris 2000).

### Microsatellites

Samples showing multi-locus matches were excluded from the dataset prior to further analysis. Expected and observed heterozygosities in each population and overall were assessed, as well as tests of significant deviations from Hardy–Weinberg equilibrium in the distribution of alleles. A likelihood-ratio test was performed to detect linkage disequilibrium between pairs of loci (Excoffier and Slatkin 1998). Obtained p-values were corrected according to Benjamini and Hochberg (1995), accounting for FDR. All loci were assessed with Microchecker (ver. 2.2.3) to assess the number of null alleles and potential allele dropout. Mean gene diversity was assessed for each population, and genetic differentiation was derived through pairwise D_est_-values for each population. Their significance was given by p-values estimated from bootstrap resampling with 1,000 permutations.

Hierarchical AMOVA testing was also applied to microsatellites, in order to depict regional clustering based on population differentiation. Different scenarios were tested according to pairwise F_st_-values obtained with the AMOVA. The genetic structure of Clark’s anemonefish was then further investigated through the program STRUCTURE (ver. 2.3.3). This method uses a maximum likelihood approach to estimate the probability of accordingly dividing all the genotypes of individuals from n populations among k clusters, k = {1,…, n}. The burn-in period was set to 100,000 iterations, and total length was set to 500,000 steps with 10 repetitions for each k-value. The distribution of the genetic clusters across populations is represented in Fig. 1d.

## RESULTS

### Neutrality tests of the genetic markers

A non-significant test of Tajima’s D failed to reject the neutrality of the mitochondrial control-region marker (Table 2, P > 0.1), thus confirming its suitability for the current analysis. Besides that, the non-significant Fu’s FS outcome provided no evidence of departure from population equilibrium, such as recent population expansion (considered significant at the 2 % level). The mismatch distribution plot (Supplementary Fig. S1) confirms this result through a decreasing curve of expected mismatch distribution. The presence of three peaks in the observed mismatch distributions may represent substructures within the dataset (Ray et al. 2003). However, both the sum of square deviations (SSD) and Harpending’s raggedness index showed no deviation from a model of sudden demographic expansion (Table 2).

**Table 2:** Neutrality tests for the suitability of mtCR marker for the current analysis of *Amphiprion clarkii*. Tajima’s D and Fu’s FS values are indicated, with corresponding p-values.

|  |  |  |
| --- | --- | --- |
| **Neutrality test** |  |  |
| Tajima's D | -1.5275 | $P > 0.1$ |
| Fu's FS | -0.9146 | $P > 0.02$ |
| **Mismatch distribution** |  |  |
| SSD | 0.0059 | $P > 0.1$ |
| Raggedness index | 0.029 | $P > 0.1$ |

Regarding microsatellites, observed heterozygosities were consistently high (Table 1, 0.924 – 1.000), each time higher than the respective expected heterozygosities. The highest number of alleles obtained was 31 for locus AC137. Little evidence of null alleles was found, the highest obtained occurrence was 24 % for CF27. No locus displayed consistent deviations from Hardy-Weinberg Equilibrium (HWE) (ChiSq > 0.05, results not shown), confirming their suitability for population genetic studies. Additionally, no population showed a consistent deviation from HWE across loci, so violation of model assuhaplotype networkions for ‘ideal’ populations is unlikely. Tests of linkage disequilibrium across pairs of loci indicate three pairs of linked loci (AC27-AC1578; AC27-AC626; 65-AC1359) across four populations (Biak, Togian Islands, Manado, Donggala). This linkage is expected to carry across populations if these markers are truly linked, which is not the case here. Therefore, all loci were expected to assort independently and were used in the following analysis.

### Genetic diversity and genetic structure

Haplotype (*h* ranging from 0.91 to 1) and nucleotide (π ranging from 1.69 to 6.35 %) diversities for the control region were consistently high among populations (Table 3). Nucleotide diversity was the highest at Biak, and overall was higher for populations located at both extremities of the archipelago. For microsatellites, observed heterozygosities were also consistently high among populations (Ho ranging from 0.81 to 0.89), being highest at Biak. Allelic richness (A) was highest at Bali and lowest at Kupang, ranging from 8.335 to 20.

**Table 3:** Sample sites, abbreviations (Abbr.), and number of individuals (Nind) per site for both markers in *Amphiprion clarkii*; number of haplotypes (Nhapl), nucleotide (π) and haplotype (h) diversity for mtCR, observed heterosigosity (Ho) and mean allelic richness (A) for polymorphic microsatellites; SD: standard deviation.

| Control Region (mtCR) |  |  |  |  |  | Microsatellites-8 loci |  |  |
| --- | --- | --- | --- | --- | --- | --- | --- | --- |
| Site | Abbr. | N<sub>ind</sub> | N<sub>hapl</sub> | *h* + SD | $\pi$ (%) + SD | N<sub>ind</sub> | Ho + SD | A + SD |
| Padang | Pa | 14 | 13 | 0.98 +/- 0.03 | 1.89 +/- 1.0 | 14 | 0.81 +/- 0.442 | 14.00 +/- 0.00 |
| Karimunjawa | Ka | 8 | 8 | 1.00 +/- 0.06 | 4.61 +/- 2.6 | 9 | 0.82 +/- 0.453 | 9.057 +/- 0.21 |
| Bali | Ba | 20 | 19 | 0.99 +/- 0.01 | 4.83 +/- 2.4 | 20 | 0.84 +/- 0.451 | 20.00 +/- 0.00 |
| Taiwan | Ta | 8 | 8 | 1.00 +/- 0.06 | 1.69 +/- 1.0 | 8 | 0.86 +/- 0.476 | 8.563 +/- 0.78 |
| Banggi Islands | Bl | 4 | 4 | 1.00 +/- 0.17 | 2.39 +/- 1.6 | 13 | 0.83 +/- 0.453 | 10.97 +/- 0.99 |
| Spermonde | Sp | 15 | 15 | 1.00 +/- 0.02 | 2.46 +/- 1.3 | 21 | 0.85 +/- 0.457 | 14.52 +/- 1.83 |
| Donggala | Do | 20 | 20 | 1.00 +/- 0.01 | 2.45 +/- 1.3 | 19 | 0.83 +/- 0.444 | 19.00 +/- 0.00 |
| Togian Islands | Tl | 9 | 9 | 1.00 +/- 0.05 | 2.59 +/- 1.4 | 17 | 0.82 +/- 0.442 | 12.43 +/- 0.96 |
| Kendari | Ke | 8 | 7 | 0.96 +/- 0.07 | 2.29 +/- 1.3 | 10 | 0.83 +/- 0.464 | 9.824 +/- 0.48 |
| Luwuk | Lu | 14 | 12 | 0.96 +/- 0.04 | 1.74 +/- 0.9 | 14 | 0.84 +/- 0.453 | 13.75 +/- 0.25 |
| Manado | Ma | 9 | 8 | 0.97 +/- 0.06 | 2.27 +/- 1.3 | 17 | 0.86 +/- 0.470 | 11.98 +/- 1.79 |
| Kupang | Ku | 10 | 9 | 0.97 +/- 0.05 | 1.91 +/- 1.0 | 10 | 0.86 +/- 0.471 | 8.335 +/- 0.34 |
| Misool | Mi | 10 | 7 | 0.91 +/- 0.07 | 3.64 +/- 2.0 | 10 | 0.86 +/- 0.472 | 10.43 +/- 1.42 |
| Pisang | Pi | 9 | 7 | 0.94 +/- 0.07 | 2.45 +/- 1.4 | 10 | 0.85 +/- 0.466 | 10.00 +/- 0.00 |
| Triton Bay | TB | 19 | 18 | 0.99 +/- 0.01 | 1.94 +/- 1.0 | 6 | 0.81 +/- 0.463 | 5.541 +/- 0.98 |
| Biak | Bk | 21 | 16 | 0.96 +/- 0.02 | 6.35 +/- 3.2 | 19 | 0.89 +/- 0.471 | 17.56 +/- 2.41 |

AMOVA performed with both markers showed highly structured populations, significantly deviating from panmictic conditions (mtCR: Φ_st_ = 0.42039, *P* < 0.0001; microsatellites: F_st_ = 0.01449, *P* < 0.0001). Pairwise Φ_st_- and D_est_-values are shown in Table 5. Both markers congruent in a very strong differentiation of Biak (pairwise Φ_st_ ranging from 0.54 to 0.65; D_est_ ranging from 0.08 to 0.1) and Padang (pairwise Φ_st_ ranging from 0.62 to 0.7; D_est_ ranging from 0.09 to 0.1). Microsatellites also supported a relatively high differentiation of Karimunjawa (pairwise D_est_ ranging from 0.07 to 0.1) from the other populations.

The best supported grouping in a hierarchical AMOVA was a differentiation between Biak [Bk] and the other populations for the control region (Table 4, Φ_ct_ = 0.63852, *P* = 0.04692), while tests performed with microsatellites rather supported the separation of Padang and Karimunjawa [Pa,Ka] additionally to Biak [Bk] from the rest of the populations (F_ct_ = 0.05199, *P* < 0.0001). Overall, both datasets distinguished populations from western and eastern IMA as being distinct from central ones.

**Table 4:** Hierarchical AMOVA groupings for different scenarios with both mtCR and microsatellites in anemonefish *Amphiprion clarkii* from the Indo-Malay Archipelago. Bold values indicate combinations providing the highest significant Φ ct value for each marker.

| # Groups | Groupings | Control region D-loop |  | Microsatellites-8 loci |  |
| --- | --- | --- | --- | --- | --- |
| | | $\Phi_{ct}$ | P | $F_{ct}$ | P |
| 2 | [Bk][all others] | **0.63852** | **0.04692** | 0.04341 | 0.06647 |
|  | [Pa][all others] | 0.30207 | 0.11241 | 0.03851 | 0.11632 |
| 3 | [Bk,Mi][Pa][all others] | 0.36895 | 0.08504 | 0.02485 | 0.02933 |
|  | [Pa][Bk][all others] | 0.62545 | 0.01369 | 0.04829 | 0.01271 |
|  | [Pa,Ka,Ba][Bk][all others] | 0.42871 | 0.01095 | 0.03001 | <0.0001 |
|  | [Pa,Ka][Bk][all others] | 0.58028 | 0.00098 | **0.05199** | **<0.0001** |
| 4 | [Bk][Mi][Ka,Pa][all others] | 0.53686 | 0.00782 | 0.04590 | <0.0001 |
|  | [Bk][Mi][Pa][all others] | 0.57371 | 0.01857 | 0.04067 | 0.00782 |
|  | [Bk,Mi][Pi][Pa,Ka][all others] | 0.38269 | 0.05474 | 0.03889 | 0.00098 |
|  | [Bk,Mi][TB][Pa][all others] | 0.25804 | 0.01003 | 0.01953 | 0.04203 |

Results given by STRUCTURE indicate the most probable scenario as being *k* = 3 clusters among the dataset (Fig. 1c; Supplementary Fig. S2), which could be attributable to the Western, Central and Eastern IMA. Indeed, the bar plot calculated with STRUCTURE shows a very clear differentiation of population 16 (Biak), and populations 1 and 2 (Padang and Karimunjawa, respectively) from the rest of the dataset (Fig. 1c). Pie charts depict the distribution of the genotype clusters identified across populations (Fig. 1d). Results are highly consistent with hierarchical AMOVA groupings performed with microsatellites (Table 4) and the haplotype network (Fig. 1a).

### Haplotype network

The haplotype network drawn from the 209 mtCR DNA sequences (Fig. 1a) showed four haplogroups (A, B, C and D), separated by 14 and 42 mutations. The most abundant one was haplogroup A, which was encountered at 14 out of 16 sample sites. It is mostly distributed throughout the central IMA, from Karimunjawa to Triton Bay. Eleven stations exclusively display this haplogroup, especially in Sulawesi where only this one was found. Due to its wide distribution and its important presence among populations, it is most likely to contain ancestral haplotypes. Haplogroup B was exclusively found at Biak. This sample site harbours haplogroups B and D, which differ by 40 and 42 mutations from haplogroup A, respectively. This makes the sample site from Biak particularly differentiated from the others. Haplogroup C was found throughout the Western IMA, from Bali to Padang. Haplogroup D is the smallest one and was only distributed in Biak and Misool. Most of the populations only harbour one haplogroup (Fig. 1b), which shows a very clear genetic structure across the IMA, and strong differentiations occurring among the Western, Central and Eastern IMA.

## DISCUSSION

### Genetic structure

The genetic breaks identified across the IMA for Clark’s anemonefish are consistent with those existing for congeneric species from the same region (*A. ocellaris*, Timm and Kochzius, 2008; Timm et al. 2012; *A. perideraion*, Dohna et al. 2015; and also several other coral reef taxa, such as giant clams (*Tridacna crocea*: Kochzius and Nuryanto 2008, Hui et al. 2017; *T. maxima:* Nuryanto and Kochzius 2009; *T. squamosa*: Hui et al. 2018), corals (*Heliofungia actiniformis*: Knittweis et al. 2009; *Seriatopora hystrix*: van der Ven et al. 2021), and a sea star (*Linckia laevigata*: Kochzius et al. 2009). The haplotype network and hierarchical AMOVA groupings provide a clear pattern of genetic differentiation among the Western, Central and Eastern IMA (Table 4, Fig. 1a). This is confirmed by the STRUCTURE analysis (Fig. 1c), as three clusters were clearly identified. Padang and Karimunjawa form the western group, and Biak alone forms the eastern group. These three sample sites also show high pairwise Φ_st_ and D_est_-values (Table 5). Overall, these groupings are also consistent with the biogeographic coastal provinces defined by Spalding et al. (2007), supporting the idea that current differentiation patterns are influenced by historical climatic events.

**Table 5:** Pairwise Φ_st_ (below diagonal) and D_es_t (above diagonal) values for the mitochondrial control region marker and microsatellites, respectively, in the clownfish Amphiprion clarkii from the Indo-Malay Archipelago. Bold values are significant (P < 0.05).

| mtCR\Msat | Mi | Pi | Pa | Ta | TB | Sp | Ka | TI | Bk | Ma | Ke | BI | Ba | Ku | Do | Lu |  |
| --- | --- | --- | --- | --- | --- | --- | --- | --- | --- | --- | --- | --- | --- | --- | --- | --- | --- |
| Mi |  | 0.046 | **0.102** | 0.055 | 0.049 | **0.048** | 0.094 | **0.059** | 0.079 | **0.056** | 0.040 | 0.054 | **0.059** | 0.033 | **0.058** | **0.052** | Mi |
| Pi | **0.02459** |  | 0.105 | **0.049** | 0.060 | 0.047 | 0.097 | 0.070 | 0.093 | 0.072 | **0.038** | 0.049 | 0.057 | 0.056 | 0.060 | 0.061 | Pi |
| Pa | 0.62796 | **0.68618** |  | 0.094 | 0.111 | **0.092** | **0.038** | **0.111** | **0.103** | 0.104 | 0.088 | **0.092** | 0.062 | **0.100** | **0.101** | 0.107 | Pa |
| Ta | **0.12941** | 0.07809 | **0.699** |  | 0.041 | 0.027 | 0.083 | **0.046** | 0.085 | **0.046** | 0.039 | 0.039 | 0.048 | 0.035 | 0.036 | **0.039** | Ta |
| TB | 0.02639 | 0.00011 | 0.69259 | 0.0843 |  | 0.026 | 0.094 | 0.051 | 0.081 | **0.042** | **0.038** | **0.033** | **0.054** | 0.038 | **0.031** | 0.040 | TB |
| Sp | **0.13907** | **0.09419** | **0.64781** | 0.00582 | 0.06942 |  | **0.080** | 0.038 | **0.076** | **0.033** | 0.028 | **0.025** | 0.042 | **0.029** | 0.028 | 0.031 | Sp |
| Ka | 0.20055 | 0.23723 | 0.31374 | 0.22803 | **0.27314** | 0.22178 |  | **0.098** | **0.102** | 0.085 | **0.074** | **0.084** | 0.057 | 0.084 | 0.083 | 0.091 | Ka |
| TI | **0.07254** | 0.01429 | 0.65645 | **0.02688** | 0.02665 | 0.5859 | 0.18234 |  | 0.102 | 0.038 | 0.039 | 0.056 | **0.062** | 0.042 | **0.050** | **0.039** | TI |
| Bk | 0.53986 | **0.59663** | **0.64612** | **0.60124** | 0.65418 | 0.62368 | **0.54213** | 0.05226 |  | **0.097** | **0.081** | 0.084 | 0.091 | 0.079 | 0.093 | **0.094** | Bk |
| Ma | 0.10104 | 0.07258 | 0.68536 | 0.09065 | **0.06432** | **0.03492** | 0.19938 | 0.0271 | 0.60784 |  | 0.036 | 0.045 | 0.053 | **0.037** | 0.036 | 0.037 | Ma |
| Ke | 0.13413 | **0.12194** | **0.65236** | **0.06286** | 0.09038 | 0.00124 | 0.17919 | 0.00841 | **0.58604** | 0.08105 |  | 0.033 | **0.040** | **0.033** | **0.033** | **0.035** | Ke |
| BI | 0.07411 | 0.08791 | 0.64207 | 0.05557 | 0.07649 | **0.0227** | 0.10373 | **0.01119** | 0.53913 | 0.03226 | **0.02487** |  | 0.042 | 0.040 | 0.037 | 0.044 | BI |
| Ba | **0.27854** | **0.28986** | 0.15918 | **0.257** | 0.32029 | 0.26342 | **0.00112** | 0.24769 | 0.56111 | **0.28118** | 0.22313 | 0.16201 |  | 0.051 | **0.049** | 0.053 | Ba |
| Ku | **0.12353** | **0.08545** | 0.67728 | 0.04055 | **0.06843** | 0.01812 | 0.2003 | -0.0074 | **0.60197** | 0.05226 | **0.0294** | -0.0161 | **0.25144** |  | 0.037 | **0.034** | Ku |
| Do | 0.10852 | 0.05355 | **0.62359** | 0.00441 | 0.03295 | -0.0038 | 0.18223 | **0.01239** | 0.63329 | **0.01045** | -0.0044 | **0.03073** | 0.245 | **0.0103** |  | **0.041** | Do |
| Lu | **0.12641** | 0.08069 | **0.69612** | **0.02683** | 0.05651 | 0.04616 | **0.24243** | -0.0071 | **0.62765** | 0.07611 | **0.01711** | 0.06728 | 0.28569 | -0.0341 | 0.00292 |  | Lu |
|  | Mi | Pi | Pa | Ta | TB | Sp | Ka | TI | Bk | Ma | Ke | BI | Ba | Ku | Do | Lu |  |

Regarding the western IMA, results support the differentiation of samples of the Eastern Indian Ocean (Padang) and Java Sea (Karimunjawa) from the central archipelago.

Hierarchical AMOVA groupings performed with microsatellites and the analysis performed with STRUCTURE group these populations together (Table 4, Fig. 1c), although disagreements with mtCR marker exist. This genetic barrier has been identified for two *Amphiprion* species (*A. ocellaris*: Timm and Kochzius 2008, Timm et al. 2012; *A. perideraion*: Dohna et al. 2015). The formation of a massive land barrier along the current Indonesian islands (Fig. 1b, light grey areas) has already been identified as a cause of allopatric speciation among populations (McMillan and Palumbi, 1995; Kochzius et al., 2003), through fluctuations of sea level and fragmentation of habitat during the Pleistocene (Voris, 2000). This study shows that this historical genetic break still exists due to a contemporary oceanographic barrier. In a biophysical modelling approach, it was shown that SSCs prevent larvae to travel from the Flores Sea to the Java Sea, causing its populations to remain isolated (Kool et al. 2011). Microsatellite markers confirm a high differentiation occurring among Western and Central populations, suggesting a continued isolation of populations of the Java Sea from the rest of the IMA. This supports the idea that old vicariant events still remain today, and act upon the genetic differentiation pattern of marine species.

Populations from the central part of the IMA only hold haplogroup A haplotypes (Fig. 1), while populations of congeneric species of the same region rather share several haplogroups (*Amphiprion ocellaris*: eight haplogroups, Timm and Kochzius, 2008; *Amphiprion perideraion*: nine haplogroups, Dohna et al. 2015). Populations of *A. ocellaris* even display a genetic differentiation between the northern (Philippines, Sulu Sea) and southern (Sulawesi, Nusa Tenggara) IMA, which is explained by a shorter PLD than its congeners (Madduppa et al. 2014). The pattern obtained for populations of Clark’s anemonefish might be due to the longer PLD of that species (Ye et al. 2011), providing a greater dispersal potential leading to a more homogenised genetic variation. This lowers the number of mutational steps between haplotypes and leads to the obtained genetic structure. This longer PLD might also explain the connectivity occurring among the central IMA and populations from western New Guinea (Triton Bay, Misool). These localities rather belong to the eastern group for congeneric species, while hierarchical AMOVA groupings and STRUCTURE analysis of Clark’s anemonefish (Table 4, Fig. 1d) show that these populations of that species rather belong to the central group.

As for the eastern IMA, the highest significant Φ_ct_-values obtained from hierarchical AMOVA groupings with both markers were obtained when Biak alone forms the eastern group. Similar groupings were identified for congeneric anemonefish species. Regarding *A. perideraion*, (Dohna et al. 2015), microsatellite data groups this population alone, while mtCR data also includes Pisang and Triton Bay as part of the eastern group. The particular genetic diversity of populations from Misool has already been mentioned (Dohna et al. 2015). The current study also provides evidence of a specific genetic diversity, as this population is the only one to harbour haplogroup D haplotypes, besides Biak (Fig. 1), although virtually no current gene flow occurs among Misool and Biak. Further studies should focus on the smallscale genetic structure of the populations located in this region. Moreover, the haplotype network and pairwise differentiation values indicate Biak as being the most distant population (Fig. 1: 40 and 42 mutations). The very high number of mutational steps among Biak and the rest of the dataset might indicate allopatric speciation occurring within this region, and the population from Biak forming a new cryptic species.

### Limited gene flow

Limited current gene flow caused local isolation and populations to evolve almost independently, shaping a strong and clear genetic structure across the IMA (Fig. 1). Gene flow is overall lower among the three different regions of the IMA (western, central and eastern part) than within regions. This overall restricted gene flow in anemonefish is commonly attributed to their short PLD (Timm and Kochzius 2008; Dohna et al. 2015; Huyghe and Kochzius, 2017; Timm et al. 2017), their high rate of self-recruitment (Madduppa et al. 2014) and oceanographic barriers (Kool et al. 2001; Huyghe and Kochzius, 2018). As an example, the strong differentiation of Biak could be explained by the formation of the Halmahera Eddy, acting as an oceanographic barrier and leading to the isolation of this population. The biophysical dispersal model simulations from Kool et al. (2011) showed evidence of limited exchange potential between this site and other populations in the IMA, and highlights the overall importance of SSC for the connectivity of marine species in this region. Moreover, the patterns obtained with simulated larvae clearly match with the direction and the intensity of SSC occurring within this region (Fig. 1b and 1d) (Kool et al. 2011).

### Conservation perspectives

Overall, the current study shows very limited gene flow among populations, recent population separation times on an evolutionary time scale and consistently high haplotype diversity and observed heterozygosity. Together these factors, as well as the strong genetic differentiation occurring between the different regions of the IMA, provide valuable information from a conservation standpoint. The existence of barriers to gene flow, as well as the persistence of the phylogeographic barriers among Western, Central and Eastern IMA, should be considered in the design of MPA networks. Very limited exchange exists among populations located at each side of the genetic breaks, thus leading to local isolation on the long run. Additionally, the high level of self-recruitment in this taxon (Beldade et al. 2012; Madduppa et al. 2014) enhances the need to act at a smaller scale, as populations may also require specific efforts on top of designing MPA networks. For instance, the population of Clark’s anemonefish from Biak is highly differentiated, and virtually no gene flow occurs with adjacent populations. According to this, the limitation of anthropogenic threats such as reef exploitation, over-harvesting and gas emissions leading to temperature increase and ocean acidification is crucial for the conservation of this genetic diversity hotspot. The conservation of host sea anemones must also be considered, as the removal of this symbionts restrains larvae settlement, thus lowering population size. Resilience of marine species after habitat degradation has already been showed (Barber et al. 2002), but one must lower anthropogenic effects at large scale to see populations replenish.

## ACKNOWLEDGMENTS

We would like to thank the Erasmus Mundus Masters Course “TROPIMUNDO” for granting a master thesis scholarship to H.D.; the German Federal Ministry of Education and Research (BMBF, Grant no. 03F0390B, 03F0472B and 03F0643B) for funding in the framework of the German-Indonesian project “Science for the Protection of Indonesian Coastal Ecosystems (SPICE)”; D. Blohm (Universität Bremen, Germany) for support and sub-project coordination; Leibniz Centre for Tropical Marine Research (Bremen, Germany) for support, cooperation, and project coordination; J. Jompa, scientists and students of the Hasanuddin University (Makassar, Indonesia) for logistics and help during field work; the competent Indonesian authorities for permits. The SPICE project was conducted and permitted under the governmental agreement between the BMBF and the Indonesian Ministry for Research and Technology (RISTEK), Indonesian Institute of Sciences (LIPI), Indonesian Ministry of Maritime Affairs and Fisheries (DKP), and Indonesian Agency for the Assessment and Application of Technology (BPPT).

## CONFLICT OF INTEREST

On behalf of all authors, the corresponding author states that there is no conflict of interest.

## SUPPLEMENTARY FIGURES

**Supplementary Figure S1:**
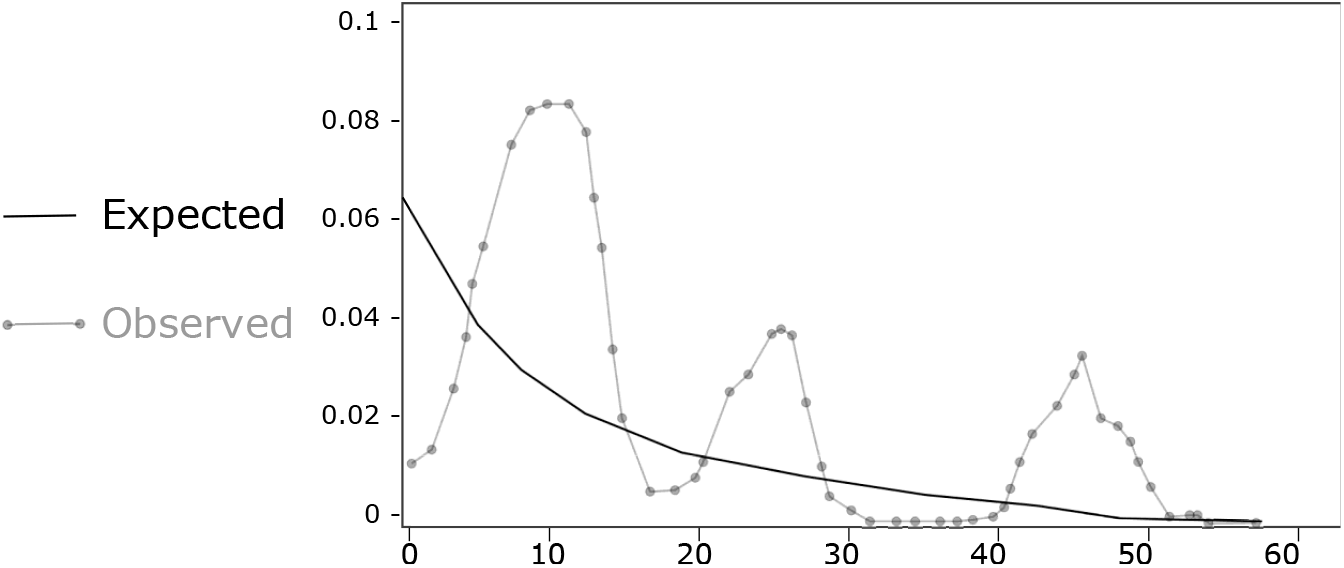
Histogram of observed versus expected frequencies of pairwise differences (mismatch distributions) for all haplotypes under a model of sudden population expansion in the anemonefish *Amphiprion clarkii* from the Indo-Malay Archipelago.

**Supplementary Figure S2:**
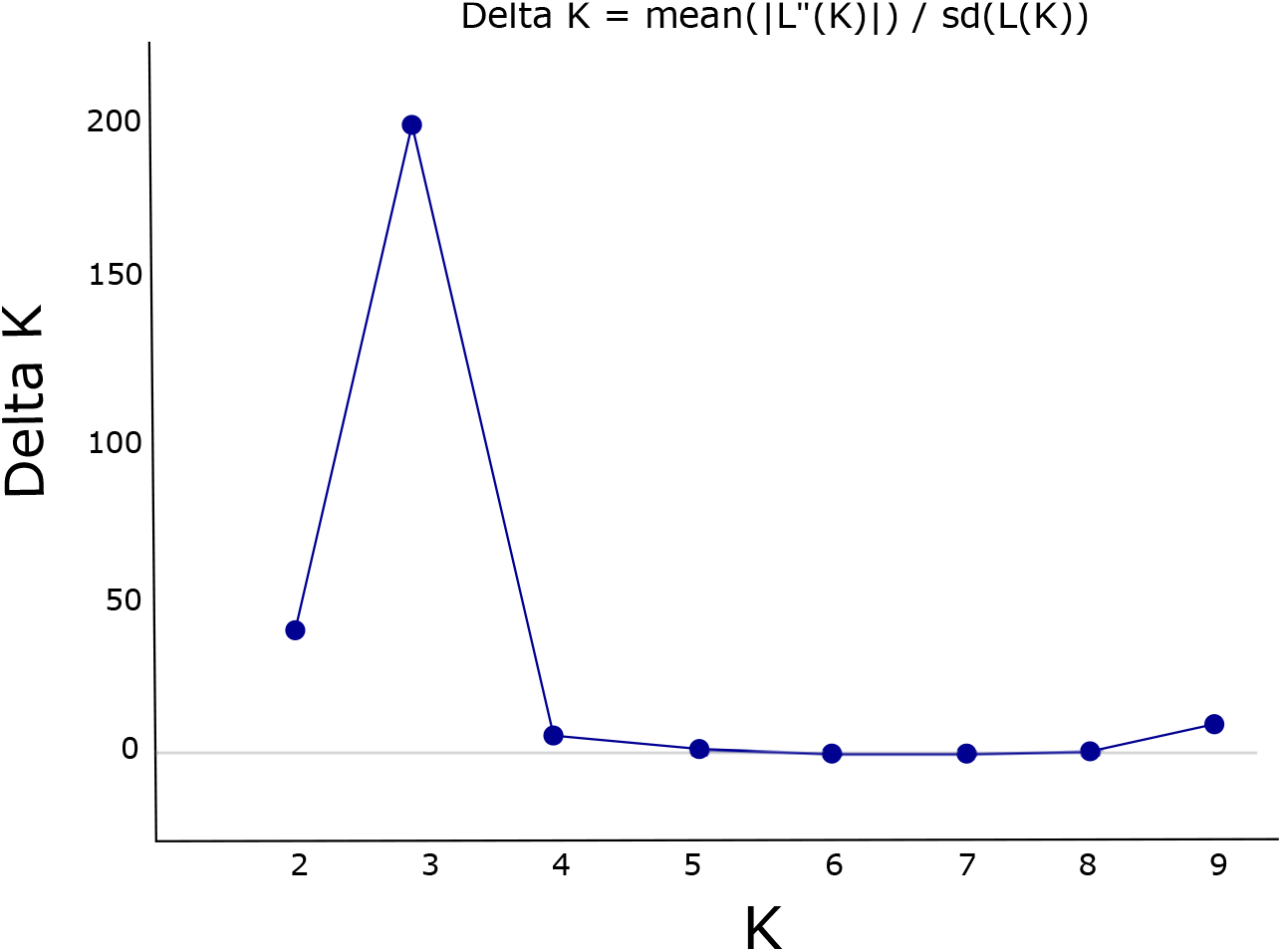
Posterior probability distribution of the Delta K curve obtained with STRUCTURE harvester. The peak of the distribution corresponds to the most probable value of k clusters in the anemonefish *Amphiprion clarkii* from the Indo-Malay Archipelago.

## Notes

### Competing Interest Statement

The authors have declared no competing interest.

